# Geographic, ecological, and temporal patterns of seabird mortality during the 2022 HPAI H5N1 outbreak on the island of Newfoundland

**DOI:** 10.1101/2024.01.17.575746

**Authors:** Gretchen M. McPhail, Sydney M. Collins, Tori V. Burt, Noah G. Careen, Parker B. Doiron, Stephanie Avery-Gomm, Tatsiana Barychka, Matthew D. English, Jolene A. Giacinti, Megan E.B. Jones, Jennifer F. Provencher, Catherine Soos, Christopher R.E. Ward, Steven Duffy, Sabina I. Wilhelm, Jordan Wight, Ishraq Rahman, Kathryn E. Hargan, Andrew S. Lang, William A. Montevecchi

## Abstract

Highly pathogenic avian influenza (HPAI) H5N1 caused mass seabird mortality across the North Atlantic in 2022. Following outbreaks in Europe, the first case in North America was detected on the island of Newfoundland (NFLD), Canada in November 2021, before spreading through all North American flyways. During the following breeding season, NFLD experienced the second-highest number of seabird mortalities in Canadian provinces. Surveys and citizen reports identified 13543 seabird mortalities from April to September 2022. Many carcasses occurred on the west coast of NFLD in May and June 2022. Reported mortalities peaked in July along the southeastern coast. In August and September, mortalities were concentrated along the northeastern coast. With the exception of two colony surveys, reported mortalities decreased in September. Most mortality was found among Northern Gannet (6622), Common Murre (5992), Atlantic Puffin (282), and Black-legged Kittiwake (217). Using comprehensive knowledge of seabird ecology, we formulated exploratory hypotheses regarding traits that could contribute to mortality. Species differences in mortality were most strongly associated with nesting density, timing of breeding, and at-sea overlap with allospecifics from other colonies. Unprecedented seabird mortality and ongoing transmission within the circulating avian influenza viruses highlight the need for continued monitoring and development of conservation strategies.

## Introduction

Seabirds are susceptible to influenza A viruses (IAVs) and their role in viral transmission across marine landscapes has been of urgent concern in recent years (Lang et al. 2016). Although seabirds are not considered a primary reservoir, diverse IAVs have been found to circulate in a wide range of seabird species, including alcids, larids, procellarids, and sphenisciformes (Ramey et al. 2010; Lebarbenchon et al. 2015; Lang et al. 2016; Barriga et al. 2016; Lee et al. 2020). Viral detections in seabirds have been primarily of the low pathogenic type of avian influenza viruses (AIVs) with a broad diversity in subtypes (Lang et al. 2016).

Beginning in 2020, highly pathogenic avian influenza virus (HPAIV) A/Goose/Guangdong/1/1996 (GsGD) H5N1 clade 2.3.4.4b caused several outbreaks in both domestic and wild bird populations across Western Europe that have continued into 2023 (European Food Safety Authority et al. 2020, 2023b). Subsequently, during the 2022 boreal summer, mass mortality in seabirds was observed across the North Atlantic Ocean as a result of HPAI H5N1 (Falchieri et al. 2022; Caliendo et al. 2022; Lane et al. 2023). In North America, a Great Black-backed Gull (*Larus marinus*) that died in November 2021 in St. John’s, Newfoundland and Labrador, Canada was infected by an H5N1 HPAIV (Caliendo et al. 2022). For several weeks prior to this, gulls with symptoms consistent with those of HPAI were reported by a local rehabilitation center (Gosse, K., pers comm. November 26, 2021). The virus was found to be closely related to viruses circulating in northwestern Europe during the spring of 2021 (Caliendo et al. 2022). This represented the first detection of HPAIV in Canada from Eurasia since an incursion in 2014 (Pasick et al. 2015). In December 2021, the virus was detected on an exhibition farm in St. John’s (Caliendo et al. 2022) and in the local wild bird population in the eastern part of the province. The first reported instance of widespread mortality in seabirds in Canada due to this virus occurred among American Common Eiders (*Somateria mollissima*) in Québec, Canada in May 2022 and was immediately followed by observations of sick and dead Northern Gannets (*Morus bassanus*) on the shores of the Gulf of St. Lawrence (Avery-Gomm et al. 2024). These first outbreaks among colonial-nesting seabirds signaled the onset of extensive mass mortality that occurred throughout eastern Canada from May to September 2022 (Giacinti et al. 2023; Avery-Gomm et al. 2024).

The island of Newfoundland (distinguished from the province of Newfoundland and Labrador, hereafter “NFLD”) was severely impacted by the HPAI H5N1 outbreak and exhibited the second highest number of mortalities, almost all of which were seabirds in Canadian provinces (Avery-Gomm et al. 2024). NFLD hosts globally significant proportions of breeding seabirds in the North Atlantic Ocean [e.g., ∼6% Northern Gannets (Chardine et al. 2013), ∼5% Common Murres (U*ria aalge*; Ainley et al. 2021), ∼6% Atlantic Puffins (*Fratercula arctica*; Lowther et al. 2020), 0.5% Black-legged Kittiwakes (*Rissa tridactyla*; Hatch et al. 2020), ∼30% Leach’s Storm-Petrels (*Hydrobates leucorhous*; Pollet et al. 2021)]. The ecology, behaviour, and movement of seabirds that breed in NFLD have been studied extensively since the 1970s (e.g., Montevecchi and Tuck 1987). To understand the local and global impacts of HPAI H5N1 on seabird species in 2022 and from potential future outbreaks, it is vital to examine the species-specific impacts that occurred during the 2022 outbreak of HPAI H5N1 in NFLD. The extensive knowledge of seabird ecology and movement in NFLD provides a compelling opportunity to investigate the roles species- and colony-specific ecology may play in possible transmission routes. It also provides the opportunity to pose exploratory hypotheses about species-specific patterns of mortality and to compare them with observed mortalities in NFLD.

Examining the traits of species that were commonly observed during the outbreak (Table 1) could help inform which traits make species more vulnerable to viral infection (van Dijk et al. 2018). As AIV transmission generally requires close contact between individuals or contact with contaminated environments (Ramey et al. 2020), we expect that species which frequently come into contact with con- and allospecifics at the colony or at sea are more likely to contract and spread the virus. Interindividual contact is likely for species that (1) nest in high densities (e.g., Common Murres, Northern Gannets; Mowbray 2020; Ainley et al. 2021), (2) interact with con- and allospecifics at the colony (e.g., Common Murres, Northern Gannets, Atlantic Puffins; Lowther et al. 2020), (3) share nesting habitat with allospecifics (e.g., cliff nesters like Black-legged Kittiwakes and Common Murres; Hatch et al. 2020; Ainley et al. 2021), and (4) overlap with con- and allospecifics at sea (e.g., Northern Gannets, Common Murres, Atlantic Puffins, Black-legged Kittiwakes; Hatch et al. 2020; Lowther et al. 2020; Ainley et al. 2021; Jeglinski et al. 2023; Careen et al. Accepted). We also hypothesize that interindividual contact is likely for species that (5) scavenge or feed on other birds (e.g., *Larus* spp.) because these species may consume infected birds and are also at risk of contracting the virus through ingestion (Brown et al. 2008). While interindividual contact is highest for most species during the breeding season, the timing of breeding varies substantially among seabird species, and some remain at the colony for much longer than others. Even within species, the timing of breeding varies among colonies, where species at higher latitudes tend to breed later (Burr et al. 2016). Depending on the regional timing of a viral outbreak, we also hypothesize that (6) some species may avoid or be more vulnerable to infection based on their timing of breeding.

**Table 1.** Seabird ecological traits that influence the level of interaction with other seabirds, predictions regarding how these traits will influence contraction, spread, and mortality from a virus (Vulnerability column), species specific traits (NOGA = Northern Gannets, COMU = Common Murre, ATPU = Atlantic Puffin, BLKI = Black-legged Kittiwake), predictions for observed mortality if the associated trait is the key factor determining infection rates, and a comparison of each prediction with the observed species-specific mortality.

| **Trait** | **Vulnerability** | **NOGA** | **COMU** | **ATPU** | **BLKI** | **Prediction** | **Result** |
| --- | --- | --- | --- | --- | --- | --- | --- |
| Nesting density | Birds that nest more densely are more likely to contact conspecifics and therefore more likely to contract and spread the virus | high | very high | moderate | moderate | COMU > NOGA >> ATPU = BLKI | Mostly Supported |
| Interaction at colony with con- and allospecifics outside of the nest site | Species which gather in groups outside of the nest at the colony (on land) are more likely to contract and spread the virus | high | moderate | high | moderate | NOGA = ATPU > COMU > BLKI | Partially supported |
| Overlap of nesting habitat among allospecifics | Species that share nesting habitat with allospecifics are more likely to contract and spread the virus | moderate | moderate | low | moderate | NOGA = COMU = BLKI > ATPU | Partially supported |
| Overlap with conspecifics from the same colony at sea | Species that interact with conspecifics at sea are more likely to contract and spread the virus | high | high | high | high | NOGA = COMU = ATPU = BLKI | Not supported |
| Overlap with allospecifics from the same colony at sea | Species that interact with allospecifics at sea are more likely to contract and spread the virus | moderate | high | high | high | COMU = ATPU = BLKI > NOGA | Not supported |
| Overlap with conspecifics from different colonies at sea | Species that interact with conspecifics at sea are more likely to contract and spread the virus | low | moderate | very low | very low | COMU > NOGA = ATPU = BLKI | Not supported |
| Overlap with allospecifics from different colonies at sea | Species that interact with allospecifics at sea are more likely to contract and spread the virus | moderate | low | very low | very low | NOGA > COMU > ATPU = BLKI | Mostly supported |
| Diet | Species that feed on other birds may contract the virus through ingestion of infected tissue; species that feed exclusively on fish, which are less likely to carry HPAI than birds, are less likely to contract the virus from ingestion of prey | very low | very low | very low | very low | Low for all | Not supported |
| Timing of breeding | Timing of breeding varies among species and colonies; also related to locations of initial outbreaks; species that breed early or have a shorter breeding season in an area where the virus reached later in the season are less likely to still be present when the virus hit their colony | April - Nov | May - Aug | May - Sept | May - Sept | NOGA > COMU = ATPU = BLKI | Mostly supported |

A wide range of birds and a number of mammalian species have been found to be infected with HPAIVs, though mortality rates depend in part on differences in taxa- and species-specific traits (European Food Safety Authority et al. 2023a; Teitelbaum et al. 2023; Alkie et al. 2023). For example, unusual mortality among dabbling ducks has not been largely attributed to HPAIV despite its detection in apparently healthy birds, though some species of diving ducks, such as Tufted Ducks (*Aythya fuligula*), are particularly susceptible to HPAIVs and often exhibit high mortality rates (Keawcharoen et al. 2008; Kleyheeg et al. 2017; Caliendo et al. 2022). Beyond host differences, factors such as viral subtype and pre-existing immunity from prior infection are important factors that affect the severity of disease due to HPAIV infection. A wide range of AIV subtypes have been identified in seabirds, with significantly varied distributions and prevalences across regions, particularly for Common Murres (Lang et al. 2016). Species and regionally specific differences in AIV prevalence and diversity thereby influence pre-existing immunity and may result in mortality differences upon infection by an HPAIV. In this study, we use the most comprehensive data available on HPAI-linked mortality in seabirds in NFLD (Avery-Gomm et al. 2024) to evaluate how species’ ecological traits are associated with susceptibility to infection and to examine exploratory hypotheses about species-specific and spatiotemporal patterns of mortality associated with the panzootic.

## Materials and methods

### Study area and data collection

We analyzed a subset of data from a comprehensive dataset of wild bird mortalities in eastern Canada from April 1 to September 30, 2022 (Avery-Gomm et al. 2024). Data included wild bird mortality observations from federal, provincial, and Indigenous governments, the Canadian Wildlife Health Cooperative, universities, social science platforms, and the public. Observations on seabird colonies were obtained by government biologists and academic researchers. The collated dataset included information on species, date, location (coordinates), observer information, and total observed mortality. Reported mortalities were attributed to HPAI if a species tested positive for the HPAI H5N1 clade 2.3.4.4b virus in eastern Canada between April 1 and September 30, 2022 (Giacinti et al. 2023). For reports with less specific taxonomic identity (e.g., unknown gull), HPAI was presumed to be the cause of mortality if more than 50% of that group of species commonly observed within the area tested positive for HPAIV. Carcasses with an unknown taxonomic identity were removed from the dataset.

In the absence of beached bird surveys and organized marking and removal of sick and dead birds from the landscape, many observations of sick and dead wild birds were opportunistic and unstructured. Observations of the same species reported two or more times within 1 km and 1 day (i.e., potentially double-counted records) were identified and removed from the dataset. A complete description of the data collation methods and processing can be found in Avery-Gomm et al. (2024). For the present study, we extracted reports of seabird mortalities (Fig. S1) in NFLD (Fig. 1) from April 1 to September 30, 2022, to focus on the seabird breeding season.

**Fig. 1.**
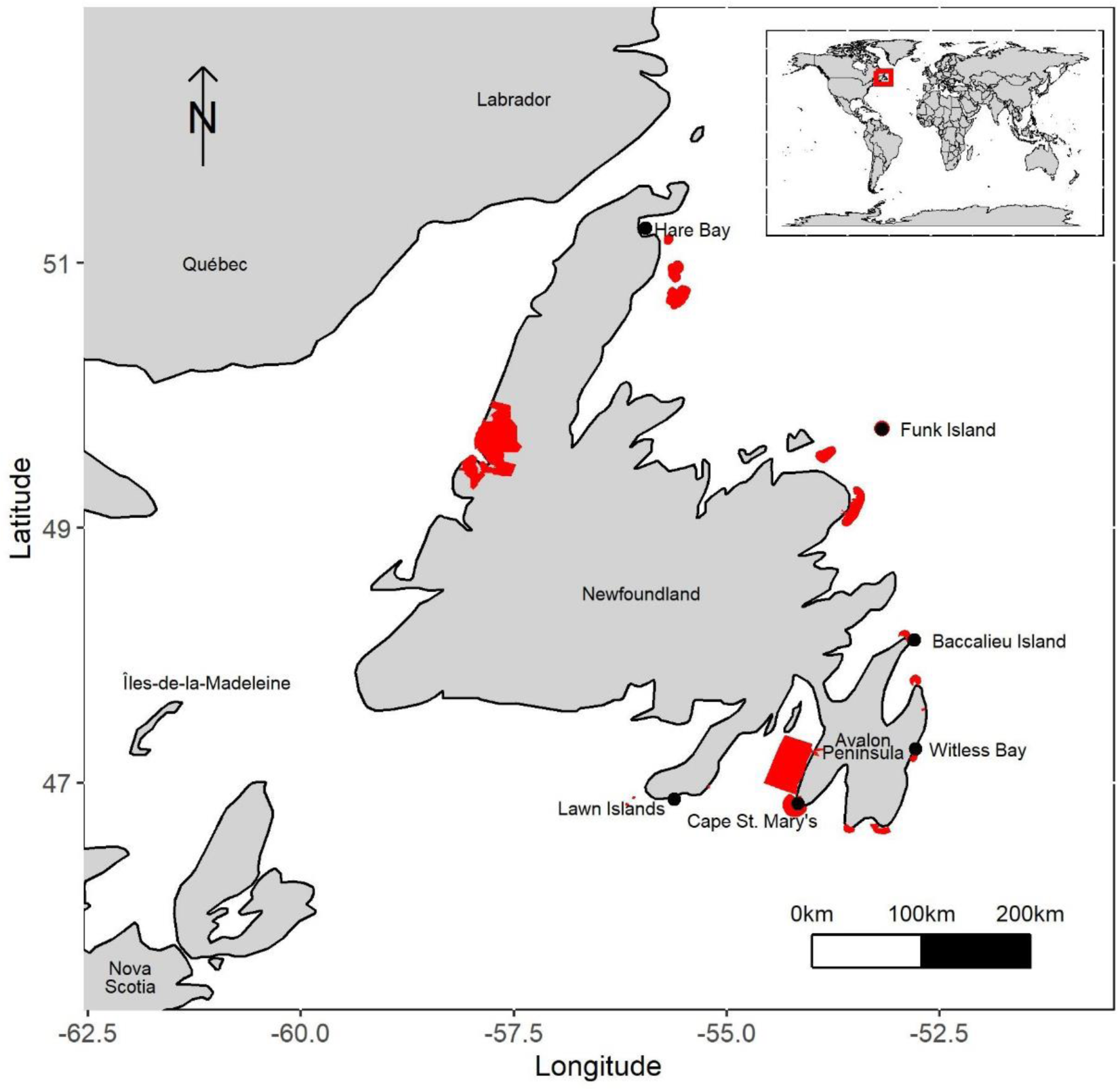
Important Bird Areas for seabirds (red polygons) (Birds Canada n.d.) and Seabird Ecological Reserves (black points) on the island of Newfoundland, Canada (Environment and Climate Change Canada 2020). Land polygons were created using the maps package (Becker et al. 2023) in R (R Core Team 2022).

### Data analysis

Analyses were conducted using R Statistical Software version 4.2.2 (R Core Team 2022). We created daily and monthly maps of the spatial and temporal patterns of mortality around NFLD. Total daily mortalities were plotted in a time-series graph to determine the period of peak mortality. To further examine spatial and temporal interactions, we plotted the daily sum of reported mortalities for the Northern and Southern areas of NFLD in relation to the colony departure chronologies of the four most impacted species. Northern regions were defined as any location north of 48.1°N latitude (the most northerly point of the Avalon Peninsula, Fig. 1). We calculated the total number of mortalities for each species and compared interspecific mortality to assess hypotheses about species traits and HPAI-associated mortality.

### Trait exploration

We created a table of seabird ecological traits which may influence the likelihood of disease transmission among con- and allospecifics from the same and different colonies (Table 1). We developed predictions for how these traits could influence transmission (and therefore the likelihood of mortality), and compared these predictions to the mortalities observed during the 2022 outbreak in NFLD.

## Results

### Spatiotemporal patterns of observed mortality

Overall, 13543 seabird mortalities were observed in NFLD from 3 April to 20 September 2022. The first eight observations of dead seabirds during the study period were made in April 2022. In May 2022, 14 seabird carcasses, primarily Northern Gannets, appeared on the west coast of the island (Fig. 2). In June, 18 mortalities were observed eastward along the south coast of the island, with four observations on the Avalon Peninsula (Fig. 2). Mortalities were concentrated along the southeast coast on the southern regions of the Burin and Avalon Peninsulas, with most mortalities (5450) observed in July (Figs 2 and 3). Mortalities continued to be high in August (3561) and September (4492, Fig. 3) and were highest on the east coast of the island (Fig. 2). Most mortalities detected in September were identified through population surveys conducted by Canadian Wildlife Service based on aerial photographs at the two largest Northern Gannet colonies in NFLD (3158 Northern Gannets at Funk Island, 1136 Northern Gannets at Cape St. Mary’s). Although these carcasses were observed and reported in September, many likely died earlier and persisted, especially at Funk Island, where the colony topography is flat, compared to the steep nesting cliffs at Cape St. Mary’s. The observed mortalities can be summarized as beginning on the west coast followed by eastward progression along the south coast, then northward along the northeast coast (Fig. S2).

**Fig. 2.**
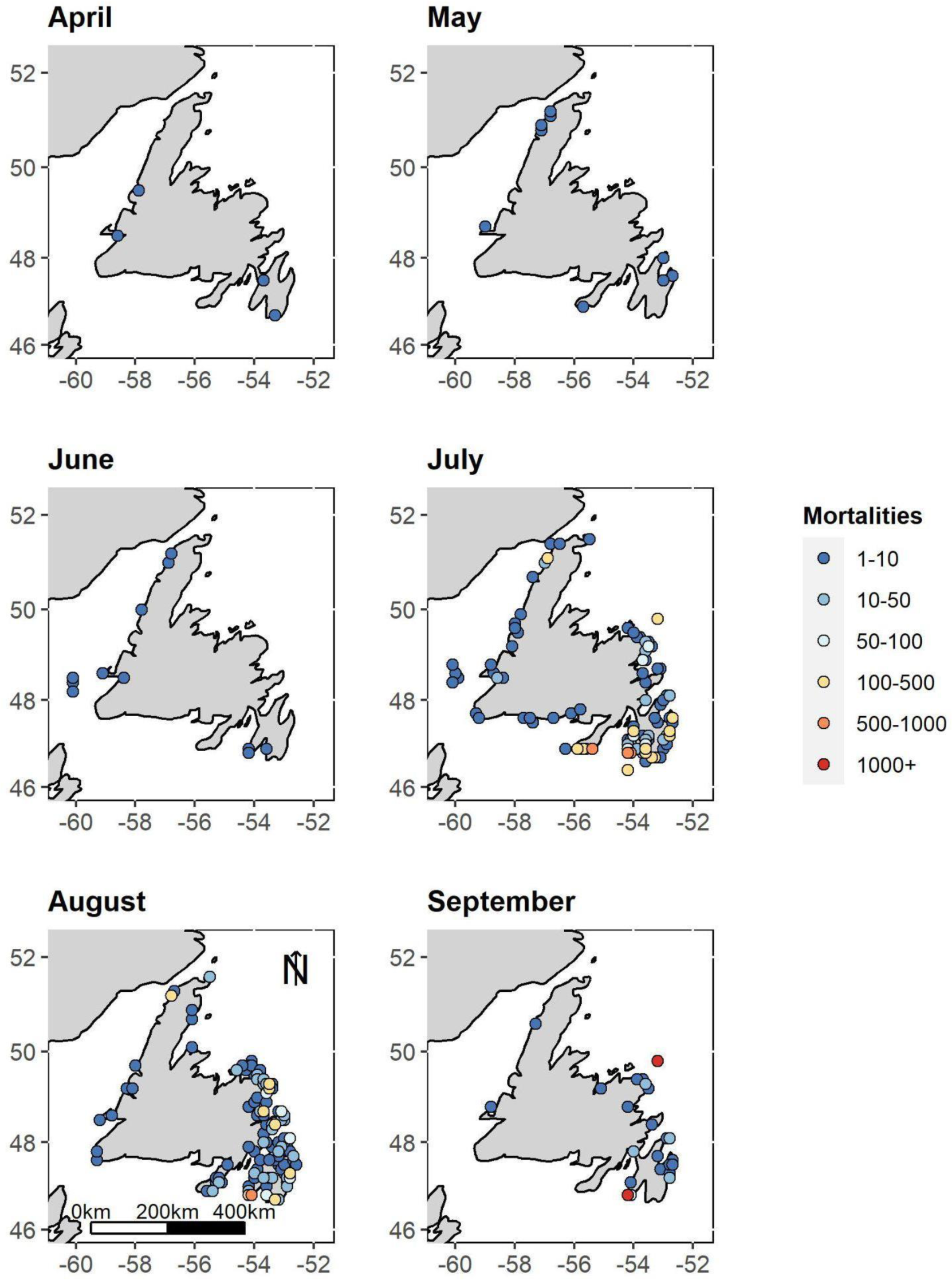
Spatial distribution of observed mortalities of seabird species along the island of Newfoundland, Canada, in April through September 2022.

**Fig. 3.**
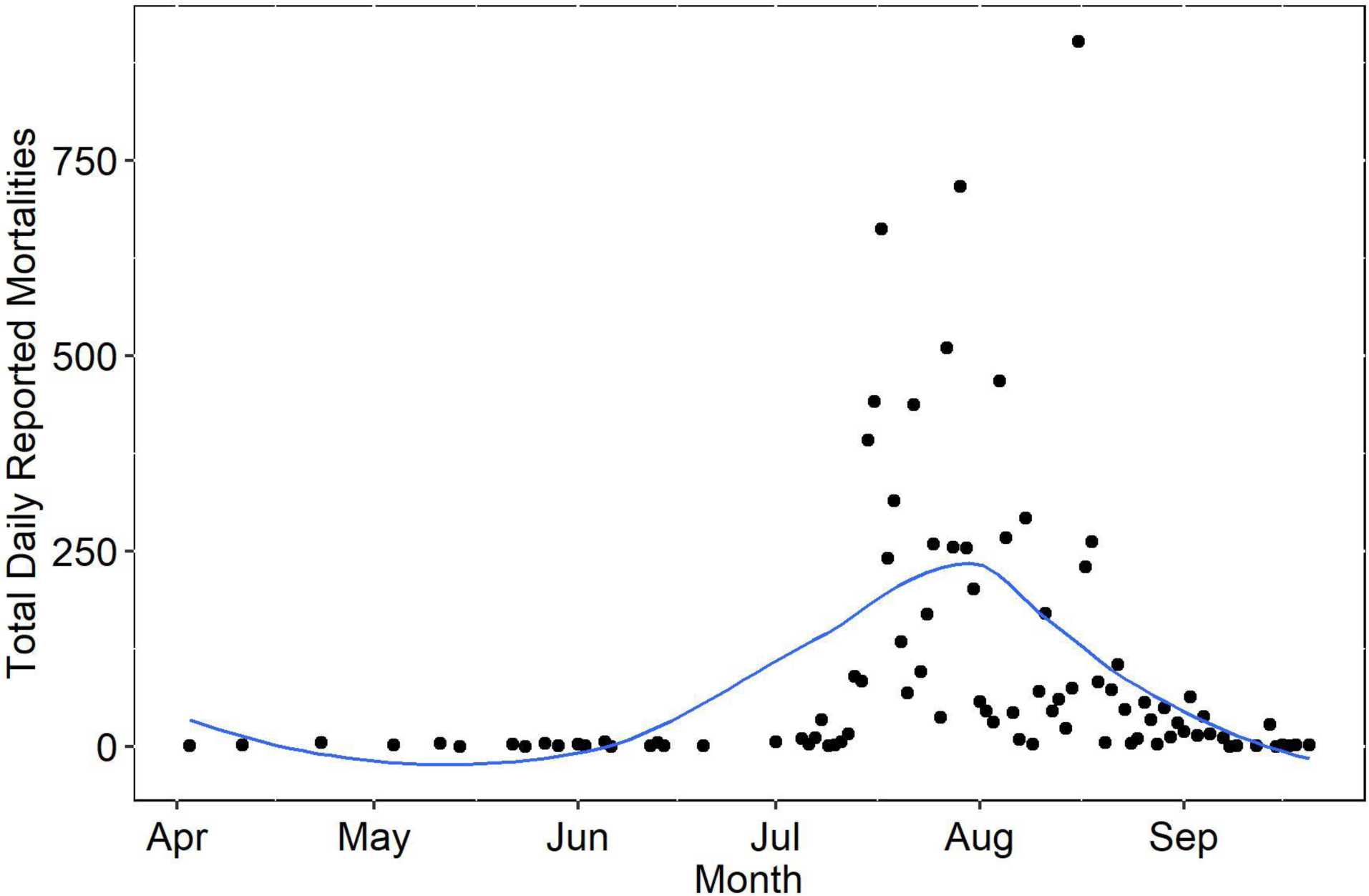
Sum of daily mortalities from 03-April to 22-Sep 2022 of seabirds on the island of Newfoundland, Canada, during an HPAI outbreak. The blue line is the LOESS line of smoothing. Note that two colony surveys conducted on 8 and 15 September 2022 were omitted from this figure, as these surveys were conducted late in the season and these dates are not representative of the dates the surveyed birds died.

### Species-specific spatiotemporal patterns of mortality

The seabird species with the highest mortalities were, in decreasing order: Northern Gannets (6622 individuals), Common Murres (5992 individuals), Atlantic Puffins (282 individuals), and Black-legged Kittiwakes (217 individuals). Razorbills (*Alca torda*) (79 individuals) and gulls also made up a large number of observed mortalities - 60 Herring Gulls (*Larus argentatus*), 31 Great Black-backed Gulls, 2 Ring-billed Gulls (*Larus delawarensis*) and 137 unidentified gulls. Notably, 27 trans-equatorial migrant Great Shearwaters (*Ardenna gravis*) were detected. Leach’s Storm-Petrels (*Hydrobates leucorhous*) were absent from the observed mortalities, despite their high regional abundance (Pollet et al. 2021).

The reported mortality of Northern Gannets and Common Murres followed a similar pattern to overall seabird mortality (Figs 4, S3-S5). Small numbers of Common Murres were first observed in early April along the west coast of NFLD but were not observed again until July (Fig. S4). In May, the first Northern Gannet mortalities were reported around the western and southern coasts, and were concentrated on the west coast in June (Fig. S5). Northern Gannet and Common Murre reports were concentrated along the southern NFLD coast in July, then along the eastern coast in August. Reported mortalities of Common Murres peaked in mid to late July (Fig. S3), whereas Northern Gannet reports peaked in early August (Fig. S3). With the exception of two colony surveys, reports of mortality for both Northern Gannets and Common Murres decreased throughout September (Figs S3-S5).

**Fig. 4.**
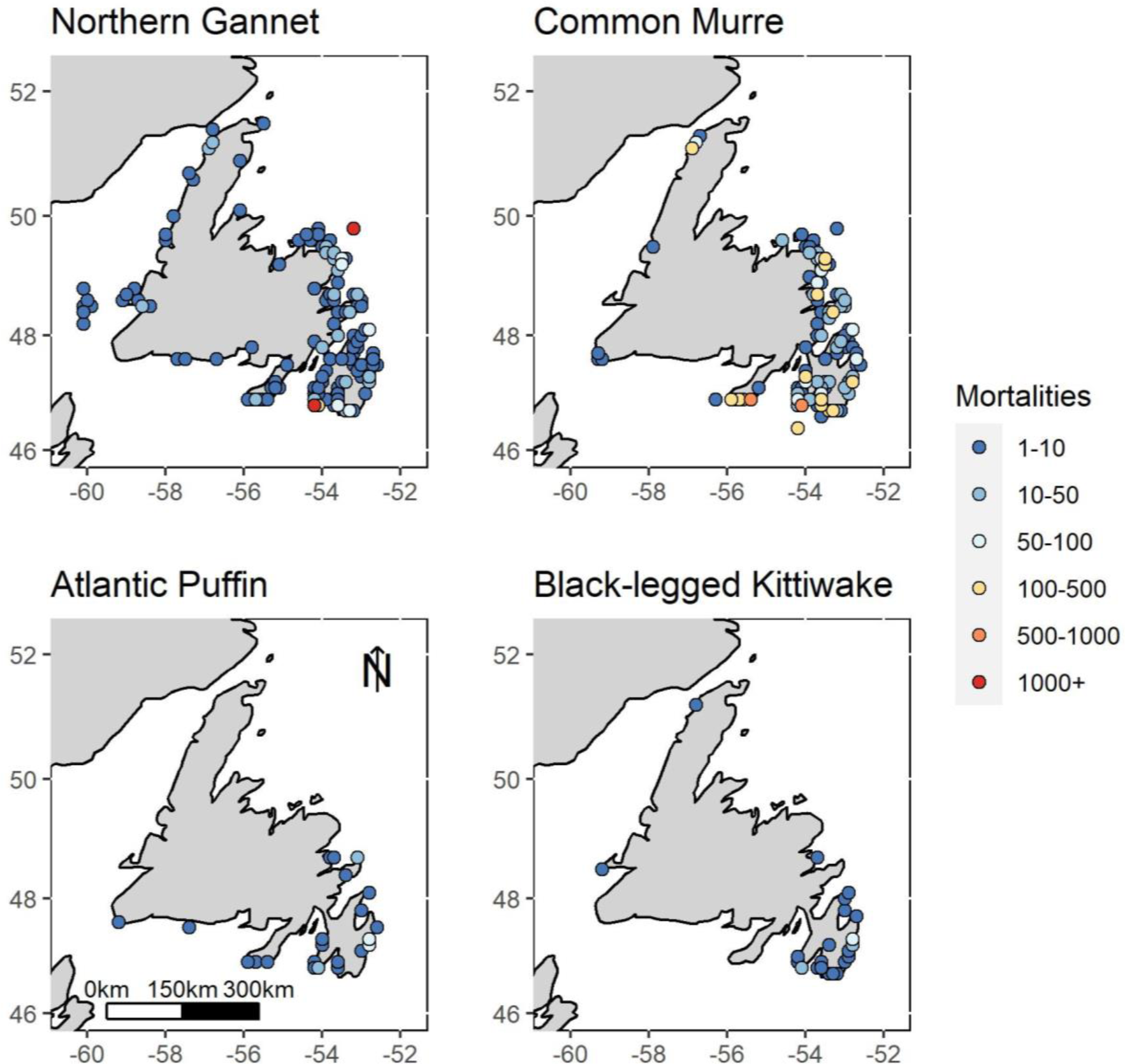
Spatial distribution of observed mortalities among Northern Gannets, Common Murres, Black-legged Kittiwakes, and Atlantic Puffins on the island of Newfoundland, Canada, from May to September 2022.

Observations of Atlantic Puffins and Black-legged Kittiwakes were first reported in July 2022 (Figs S3, S6, S7). Black-legged Kittiwake reports were concentrated almost exclusively in eastern NFLD on the Avalon Peninsula throughout the breeding season (Fig. S6). In July, Atlantic Puffin mortalities occurred along the southern coast and in August, they were observed northward along the east coast (Fig. S7). Both Atlantic Puffin and Black-legged Kittiwake mortalities decreased throughout September (Figs S3, S6, S7).

### Trait analysis

Three seabird ecological traits were most closely associated with the species-specific observed mortality in NFLD. 1) Nesting density likely has an important association with species differences in observed mortality; Northern Gannets and Common Murres have the highest nesting densities of seabirds that breed in NFLD (Table 1), and they experienced observed mortalities that were an order of magnitude greater than all other species (Fig. S1). 2) The at-sea overlap with allospecifics from different colonies was also likely associated with observed mortality, and may also relate to inter-colony transmission of the virus. 3) The timing of breeding was an important attribute with important interactions with other traits; Northern Gannets, which have the longest breeding season, experienced greater mortality than other seabirds with shorter breeding seasons (Table 1).

Allospecific overlap of nesting habitat partially predicted observed mortalities, as Northern Gannets and Common Murres tend to nest close together. However, Black-legged Kittiwakes also tend to share nesting habitat with Northern Gannets and Common Murres, yet Black-legged Kittiwakes were the fourth most observed species. Interactions at the colony with con- and allospecifics outside of the nest also partially predicted observed mortality, as Northern Gannets and Atlantic Puffins both roost or stand on cliffs next to their nests, yet there was a large disparity in observed mortality between these species.

Four seabird ecology traits were not associated with observed mortality. The predictions made with respect to species diet, overlap with con- and allospecifics from the same colony at sea, and overlap with conspecifics from different colonies at sea may have interacted with other traits, but were likely not drivers of species differences in observed mortality.

## DISCUSSION

### Seabird mortality and spatiotemporal patterns

During the 2022 HPAI H5N1 panzootic, 13543 sick and dead seabirds were reported in NFLD. Northern Gannets, Common Murres, Atlantic Puffins, and Black-legged Kittiwakes had the highest mortalities (Fig. S1). We hypothesized that species ecology might help explain interspecies differences in the incidence of infection or mortality from the virus.

Both within NFLD and throughout their breeding range, Northern Gannets experienced substantial impacts from the HPAI H5N1 panzootic (Lane et al. 2023; Wilhelm et al. in prep.). Northern Gannets began washing ashore on the west coast of the island in May-June 2022, and drift modeling suggests that these birds they likely originated from the Rochers aux Oiseaux colony in the Îles-de-la-Madeleine in the province of Québec (Avery-Gomm et al. 2023). After these first mortality reports, seabird mortalities were observed with increasing frequency along the southern NFLD coast in July, then along the east and northeast coasts in August.

### Mortality estimate considerations

Our results should be viewed as minimal mortality estimates as it is impossible to account for every bird that died from infection of HPAI. Birds that die at-sea and do not reach shore are essentially not detected and thereby grossly reduce mortality estimates. To account for these missing data is a complicated exercise, though efforts attempt to bridge this knowledge gap. For example, experimental studies of releases of marked birds (and/or decoys) at sea when systematic beach surveys were ongoing indicate that only a small proportion of birds that die at sea are found on beaches (Wiese and Robertson 2004; Munilla et al. 2011). Expansion or inflation factors based on studies like these have been used to account for losses of murres in the Pacific and Atlantic Oceans (e.g., Wiese 2002; Wiese and Robertson 2004; Piatt et al. 2020). These estimates are likely a closer approximation of true losses, however, because environmental conditions, distance from the coast, and carcass sinking greatly affect deposition on beaches, a comprehensive correction factor has yet to be derived to account for mortality at sea. Hence, actual mortality is at least several times higher than observed mortality, regardless of species (Wiese et al. 2001; Wiese 2002). Beached bird surveys are useful to detect mass mortality events and to obtain reliable information on species composition as well as age and sex, and with appropriate correction factors, to quantify to some extent mortalities associated with various threats, but not to estimate population-level impact, as more research is needed to account for species-specific carcass losses at sea (Jones et al. 2023).

Limitations and biases of citizen science data also influence observed mortality reports. In addition to the lack of reporting at sea, citizen science reports also have inherent spatial and temporal biases. Mortality reports were concentrated in areas with high human and avian population densities (e.g., the Avalon Peninsula; Fig. 2). The concentration of observations in these areas could result from a combination of the high numbers of breeding seabirds and increased detection probabilities associated with the greater human population (e.g., Burt et al. 2023). Temporal bias is also likely, as summer facilitated more participation in outdoor activities, increasing the likelihood of finding sick and dead birds. Despite these limitations of citizen science reports, the information provided important qualitative and quantitative data that would otherwise be inaccessible to research scientists (Burt et al. 2023).

From these data, we determined species-specific spatiotemporal patterns of observed mortality. By investigating the ecological traits of the most impacted species, we can make inferences regarding which traits could influence a species’ likelihood of infection (but not necessarily mortality) (Table 1). However, as mentioned, previous exposure to AIVs and immunity from prior infection also play a role in the likelihood of infection and mortality from HPAI infection, in addition to host differences.

### Associations between species ecology and likelihood of infection

#### Interactions with conspecifics and allospecifics at the colony

The two species with the highest number of observed mortalities, Northern Gannets and Common Murres, are abundant in NFLD [tens or hundreds of thousands, respectively (Chardine et al. 2013; Ainley et al. 2021)] and also have the highest nesting densities, suggesting that inter-nest distance is positively associated with the likelihood of infection (Fig. S8). Common Murres often breed shoulder-to-shoulder, and Northern Gannets nest in close proximity that allows contact between neighbouring pairs (Nelson 1978; Falchieri et al. 2022). High breeding densities in Common Murre and Northern Gannet colonies create favourable conditions for viral transmission directly from other infected birds, carcasses, aerosols, and environmental (fecal or water) contamination (Boulinier 2023).

Atlantic Puffins and Black-legged Kittiwakes also nest in large colonies in NFLD (Hatch et al. 2020; Lowther et al. 2020), though they nest further apart than Northern Gannets and Common Murres. Atlantic Puffins nest in burrows and do not contact conspecifics other than their mate while incubating, and although it is possible birds died in the burrow and were undetected, the lack of observations of this despite extensive puffin burrow monitoring across NFLD in 2022 indicates that this is unlikely (R. Zabala Belenguer pers. comm. January 2, 2024). Black-legged Kittiwakes nest on steep, sloping and sheer cliff ledges (Hatch et al. 2020) and unlike Common Murres and Northern Gannets, often have more space between nests that restrict contact between neighbouring birds. Similarly, Razorbills, the species with the fifth highest observed mortality (excluding unidentified gulls and alcids), nest in rock crevasses on cliffs with nesting densities similar to those of Black-legged Kittiwakes and Atlantic Puffins (Lavers et al. 2020). The high number of observed mortalities of Northern Gannets and Common Murres compared to the relatively low observed mortalities of Atlantic Puffins and Black-legged Kittiwakes suggest that species-specific nesting density and likelihood of infection are positively related (Fig. S8).

Sharing nesting habitat may also explain why dead Black-legged Kittiwakes, Northern Gannets, Razorbills and Common Murres were commonly observed. These species nest next to or above one another on cliffs at colony perimeters and often make direct contact with one another or con- and allospecific feces, which might result in interspecies viral transmission within a colony. Northern Gannets, Atlantic Puffins, Black-legged Kittiwakes, Common Murres, and Razorbills also interact outside of the nest while resting on land (e.g., ponds, coastal boulders, cliffs without nests) (SMC pers. obs.), and puffins interact socially outside burrows (Creelman and Storey 1991). These interactions provide further opportunities for viral spread among con- and allospecifics within a colony. These interactions, however, are far less frequent than between densely nesting neighbours, and would thus be a less significant driver of species-specific infection and mortality than nesting density.

#### Interactions with conspecifics and allospecifics at sea

While at sea, individuals could make contact with conspecifics and allospecifics from the same colony, which is most likely near a large dense multispecies colony. The foraging ranges of Atlantic Puffins, Common Murres, and Black-legged Kittiwakes overlap with other individuals from the same colony (Wanless et al. 1990; Gulka et al. 2020; Petalas et al. 2021), and these species rest together on the water in very large groups. These at-sea interactions are another possible opportunity for intra-colony viral transmission. In contrast, Northern Gannets have a much larger foraging range than Atlantic Puffins, Black-legged Kittiwakes, and Common Murres (d’Entremont et al. 2022), so are less likely to overlap at sea with allospecifics from the same colony, but forage and raft with conspecifics at foraging hotspots (e.g., Carter et al. 2016; d’Entremont et al. 2022). All four species feed on capelin when it is available early in the breeding season (Hatch et al. 2020; Lowther et al. 2020; Ainley et al. 2021; d’Entremont et al. 2022), so allospecific interaction at sea with individuals from the same colony is likely during this time. When capelin complete spawning, gannets forage farther away from the breeding colony for larger fish like mackerel (d’Entremont et al. 2022), and thus are less likely to share foraging areas with the smaller seabirds later in the breeding season.

Potential inter-colony interactions of infected individuals may also be identified by foraging ranges and at-sea space use. Atlantic Puffins tend to forage within 5 km of their breeding colony (Lowther et al. 2020), and Common Murres and Black-legged Kittiwakes stay within 30-60 km (Hatch et al. 2020; Ainley et al. 2021). The foraging ranges of many seabirds tend to scale with colony size (Patterson et al. 2022), and Common Murres from Funk Island are known to overlap at sea with murres from the small nearby (ca. 10000 breeding pairs; 66 km; Wilhelm et al. 2015) colony at South Cabot Island (Gulka et al. 2020), but the geographic separation between large seabird colonies in NFLD reduces the likelihood that Atlantic Puffins, Black-legged Kittiwakes, or Common Murres overlap with seabirds from other large colonies. Northern Gannets, however, often travel hundreds of kilometres on a single foraging trip (Mowbray 2020; d’Entremont et al. 2022) and may be more likely to overlap with individuals from other colonies in locations. For example, Northern Gannets from Cape St. Mary’s often forage in Witless Bay (d’Entremont et al. 2022), overlapping with Atlantic Puffins, Common Murres, and Black-legged Kittiwakes breeding at those colonies. However, overlap at sea with conspecifics from other colonies is unlikely for Northern Gannets, as they tend to exhibit at-sea ecological segregation among colonies (Wakefield et al. 2013). Tracking data for Northern Gannets, Common Murres, Black-legged Kittiwakes, and Atlantic Puffins are not available for all of their major colonies in NFLD, so it is difficult to verify the degree to which these species overlap with con- and allospecifics from other NFLD colonies. Continued widespread tracking of these species will help establish possible shared foraging hotspots in NFLD that may be locations where the virus is spread among individuals of different colonies.

In rare instances, Northern Gannets have also been observed visiting other colonies (Jeglinski et al. 2023; Careen et al. Accepted). Inter-colony movements by a gannet breeding at Cape St. Mary’s to Baccalieu Island were tracked during the 2022 HPAI outbreak (Careen et al. Accepted). The outbreak at Cape St. Mary’s occurred before the one at Baccalieu Island (Fig. S2), and such intercolony movement may have facilitated the viral spread to Baccalieu and other colonies. Inter-colony movements by Northern Gannets were also implicated in viral spread between colonies in Scotland during the European HPAI H5N1 outbreak (Jeglinski et al. 2023). Northern Gannets may, therefore, play a key role in the transmission of HPAI H5N1 between seabird colonies in the North Atlantic (Jeglinski et al. 2023; Careen et al. Accepted).

Great Shearwaters are common in the waters around NFLD seabird colonies (Carvalho and Davoren 2019) and were among the observed mortalities in NFLD during the H5N1 outbreak. Great Shearwaters form rafts of dozens to hundreds of individuals near shore (Rowan 1952; Brooke 1988) and could have been exposed to the virus through contact with infected allospecifics. These shearwaters are trans-equatorial migrants (Schoombie et al. 2018) and can travel many hundreds kilometers per day, with the potential to carry HPAIV across large geographic areas.

#### Diet

The majority of seabirds, including Northern Gannets, Common Murres, Black-legged Kittiwakes, and Atlantic Puffins, feed almost exclusively on fish and marine crustaceans (Hatch et al. 2020; Lowther et al. 2020; Mowbray 2020; Ainley et al. 2021), species not known to be infected by AIVs. Therefore, diet is an unlikely source of infection for primarily piscivorous seabirds.

On the other hand, *Larus* gulls (e.g., Great Black-backed Gulls, Herring Gulls, Ring-billed Gulls) are omnivores that commonly prey on other birds or scavenge carcasses (Good 2020; Weseloh et al. 2020). Many dead gulls presumed to be infected with HPAIV were reported in NFLD between April and September 2022. These gulls may have become infected by consuming infected carcasses (Brown et al. 2008). Gulls, however, are known to be frequently infected by diverse AIVs (Wille et al. 2011; Huang et al. 2014; Benkaroun et al. 2016), with previous exposure and pre-existing immunity playing an important role on their outcomes from HPAI H5N1 infection (Tarasiuk et al. 2022), therefore inferences regarding mortality among gulls were not examined in the present study.

#### Timing of Breeding

Fledging and subsequent colony departure of Atlantic Puffins, Black-legged Kittiwakes, and Common Murres peaks in early to mid-August for colonies in southern Newfoundland and Labrador (Tranquilla 2014; Hatch et al. 2020; Wilhelm et al. 2021). The fledging and departure period of Common Murres, Atlantic Puffins, and Black-legged Kittiwakes is slightly later (mid-late August to early September) for more northerly colonies (Tranquilla 2014; Hatch et al. 2020; Zabala Belenguer 2023). Birds are less likely to interact during migration when individuals are more dispersed, and based on their breeding chronology, some Common Murres, Black-legged Kittiwakes, and Atlantic Puffins may have begun migration before the viral outbreak at their colony and avoided infection (Fig. S9). Additionally, birds which were infected shortly before departure may have died at sea where their carcasses were unlikely to be observed in the colony surveys.

In contrast, gannets that breed in North America generally depart colonies from early September to mid-October (Fifield et al. 2014). Unlike kittiwakes and puffins, which leave their chicks unattended while foraging, gannet chicks are usually attended by at least one parent throughout the chick-rearing period (Lewis et al. 2004) and so are more likely to die at the nest where they were observed in colony surveys. Because gannets are present in high densities at the colony for an extended period, they are likely to contact conspecifics at the colony during a viral outbreak, regardless of the timing of the outbreak (Fig. S9). The duration of the breeding season is likely an important factor which influenced the number of observed mortalities during the HPAI H5N1 outbreak in NFLD.

#### Leach’s Storm-Petrels: An interesting exception

Interestingly, Leach’s Storm-Petrels, the most abundant procellariiform in the North Atlantic and the most abundant breeding seabird in eastern Canada (Pollet et al. 2021), were seemingly unaffected by HPAI H5N1 in 2022. Although hundreds of dead Leach’s Storm-Petrels were found and sampled during the viral outbreak, no live or dead storm-petrels sampled in 2022 were AIV positive (Avery-Gomm et al. 2024, J. Wight, unpubl. data.). Storm-petrels are attracted to anthropogenic light, which is a major source of mortality for this species, and their carcasses are commonly observed around brightly lit industrial sites (Wilhelm et al. 2021; Burt et al. 2023). Alternate causes of mortality like this emphasize the importance of viral testing to confirm the cause of death during disease outbreaks.

Storm-petrel ecology may partially explain their seeming lack of infection. They forage hundreds of kilometres offshore beyond the foraging ranges of gannets, auks, and gulls that breed in NFLD, and are unlikely to contact infected allospecific birds at sea (Ronconi et al. 2022). They also nest in burrows, which generally restrict contact with conspecifics, and are nocturnal (Pollet et al. 2021), so they are less likely to contact allospecific birds at the colony. On the other hand, storm-petrels in NFLD breed in extremely large colonies of hundreds of thousands to millions of birds (Pollet et al. 2021), and contact among conspecifics does occur, as they occasionally collide in the air (SMC pers. observation), enter neighbouring burrows (Pollet et al. 2021), or contact feces at the colony. There is also some foraging range overlap among NFLD colonies (Hedd et al. 2018) and interindividual foraging range overlap within a colony (Collins et al. 2022). It is especially surprising that the virus was not detected in storm-petrels as other procellariiforms that breed in very low numbers in NFLD, (i.e., Northern Fulmars *Fulmarus glacialis*; Garthe et al. 2004) and do not breed in NFLD but are common in coastal waters (e.g., Great Shearwaters; Carvalho and Davren 2019) were among the reported mortalities and positive for HPAIV.

### Future Research Considerations

The HPAI H5N1 panzootic has resulted in the deaths of tens of thousands of wild birds globally and poses a continued threat to domestic poultry operations (European Food Safety Authority et al. 2023a; Lane et al. 2023; Avery-Gomm et al. 2024). Due to the high burden of HPAI in wild birds globally, these viruses have been increasingly found to infect non-bird taxa including marine mammals, wild canids, and bears (Jakobek et al. 2023; Puryear et al. 2023; Alkie et al. 2023).

Wildlife managers, researchers, and the public can adopt measures to aid understanding of the impact on seabird populations following the 2022 HPAI outbreak. During a viral outbreak, standardized systematic surveys on and closely surrounding seabird colonies are needed to more precisely estimate infection-related mortality, and are important to conduct following an outbreak to estimate actual population losses (Cunningham et al. 2022). Encouraging the public to report out-of-the-ordinary occurrences regarding seabirds enables monitoring over a much larger geographic and temporal scale than is achievable through research surveys. Educating the public about species identification and behaviour will also enhance the utility and accuracy of citizen science reporting.

As HPAIVs continue to circulate in wild bird populations around the globe with possible reinfection of these seabird species and populations, it will be highly informative to assess the behavioural ecology and vital rates (reproductive success, recruitment survival) of the surviving birds to assess longer-term effects beyond immediate mortality. Using behavioural ecology to understand species’ risks and the possible routes of viral transmission can aid government wildlife and health agencies in assessing risk and potential population-level impacts from future HPAI panzootic events affecting wild birds.

## Acknowledgements

This paper is developed from the undergraduate thesis of GMM. This dataset is largely made of reports from members of the public, and we extend our sincere thanks to those who contributed reports on the island of Newfoundland. Thank you to Kyle d’Entremont and Raul Zabala Belenguer for sharing their knowledge of Newfoundland seabird ecology.

## Author contribution statement

**GMM**: Conceptualization, Data curation, Formal Analysis, Investigation, Methodology, Visualization, Writing – original draft, Writing - review & editing; **SMC**: Conceptualization, Data curation, Formal Analysis, Investigation, Methodology, Supervision, Visualization, Writing – original draft, Writing - review & editing; Supervision; **TVB**: Conceptualization, Data curation, Investigation, Methodology, Writing – original draft, Writing – review & editing; **NGC**: Investigation, Writing – review & editing; **PBD**: Investigation, Methodology, Writing – original draft, Writing - review & editing; **SA-G**: Conceptualization, Data curation, Formal analysis, Investigation, Methodology, Writing - review & editing; **TB**: Data curation, Formal analysis, Investigation, Methodology, Visualization, Writing - review & editing; **MDE**: Data curation, Writing - review & editing; **JAG**: Investigation, Writing - review & editing; **MEBJ**: Investigation; **JFP**: Investigation, Writing - review & editing; **CS**: Investigation; **CREW**: Investigation, Writing - review & editing; **SD**: Investigation, Writing - review & editing; **SIW**: Investigation, Writing - review & editing; **JW**: Conceptualization, Data curation, Investigation, Methodology, Writing – original draft, Writing – review & editing; **IR**: Data curation, Investigation, Methodology, Writing – review & editing; **KEH**: Supervision; **ASL**: Conceptualization, Data curation, Funding acquisition, Investigation, Methodology, Writing – original draft, Writing – review & editing, Supervision; **WAM**: Conceptualization, Data curation, Funding acquisition, Investigation, Methodology, Visualization, Writing – original draft, Writing – review & editing, Supervision.

## Data Availability

Data analyzed during this study (Data S1) will be available upon publication on FigShare at the following doi: https://doi.org/10.6084/m9.figshare.24856869

