## Supplementary Material for "Geographic, ecological, and temporal patterns of seabird mortality during the 2022 HPAI H5N1 outbreak on the island of Newfoundland"

### Supplementary Figures

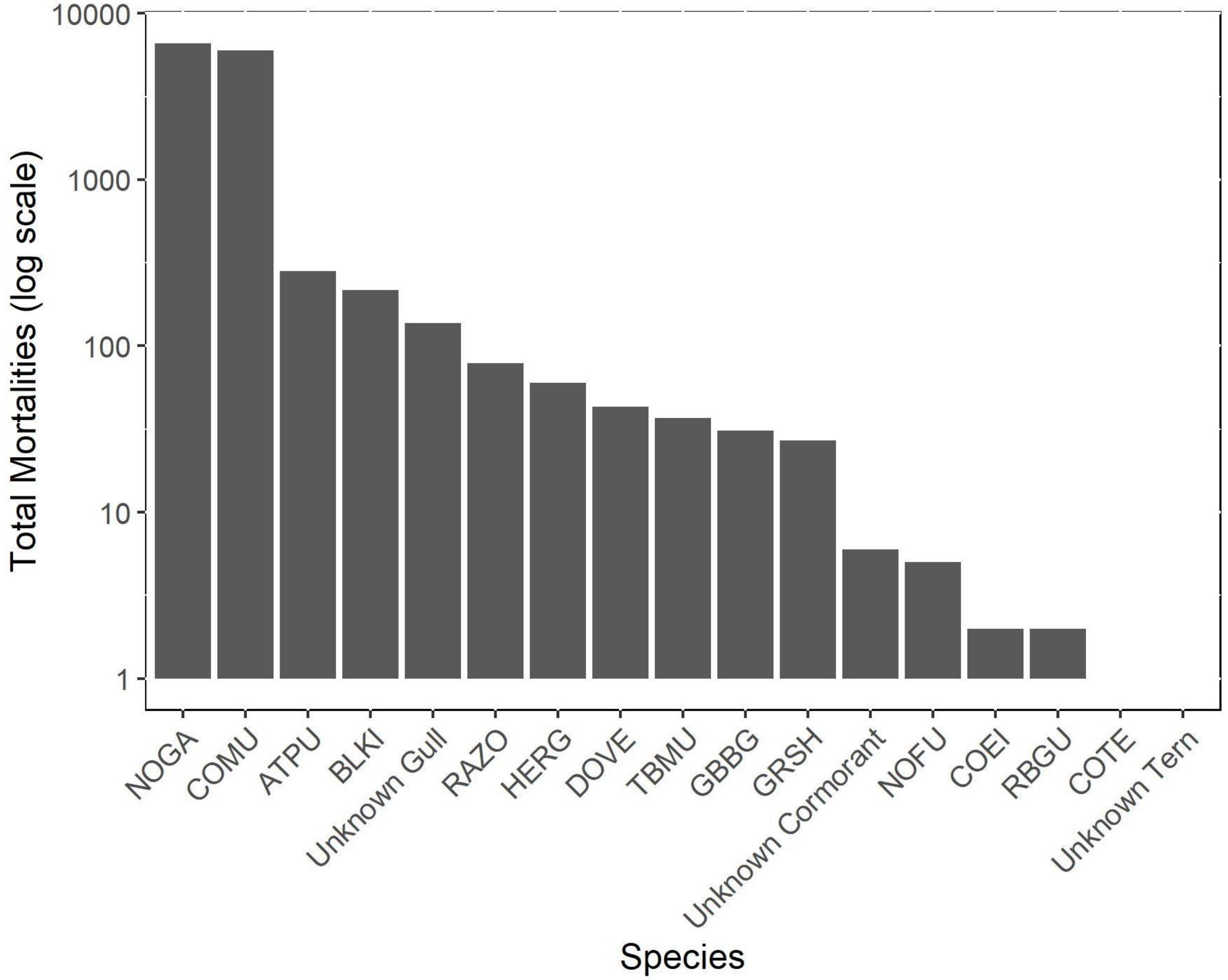


Fig. S1. Mortalities reported per species using alpha code nomenclature (https://www.pwrc.usgs.gov/BBL/Bander_Portal/login/speclist.php), plotted on a logarithmic scale.

*See multimedia file “McPhail_etal_2024_NFLD_HPAI.gif”*

Fig. S2. Video of the observed mortalities of seabirds across the island of Newfoundland from April to September 2022. Note the initial outbreak on the west coast, followed by the progression to the southeastern coast, then progression northward along the east coast. Observations made on the same day at the same location were summed.

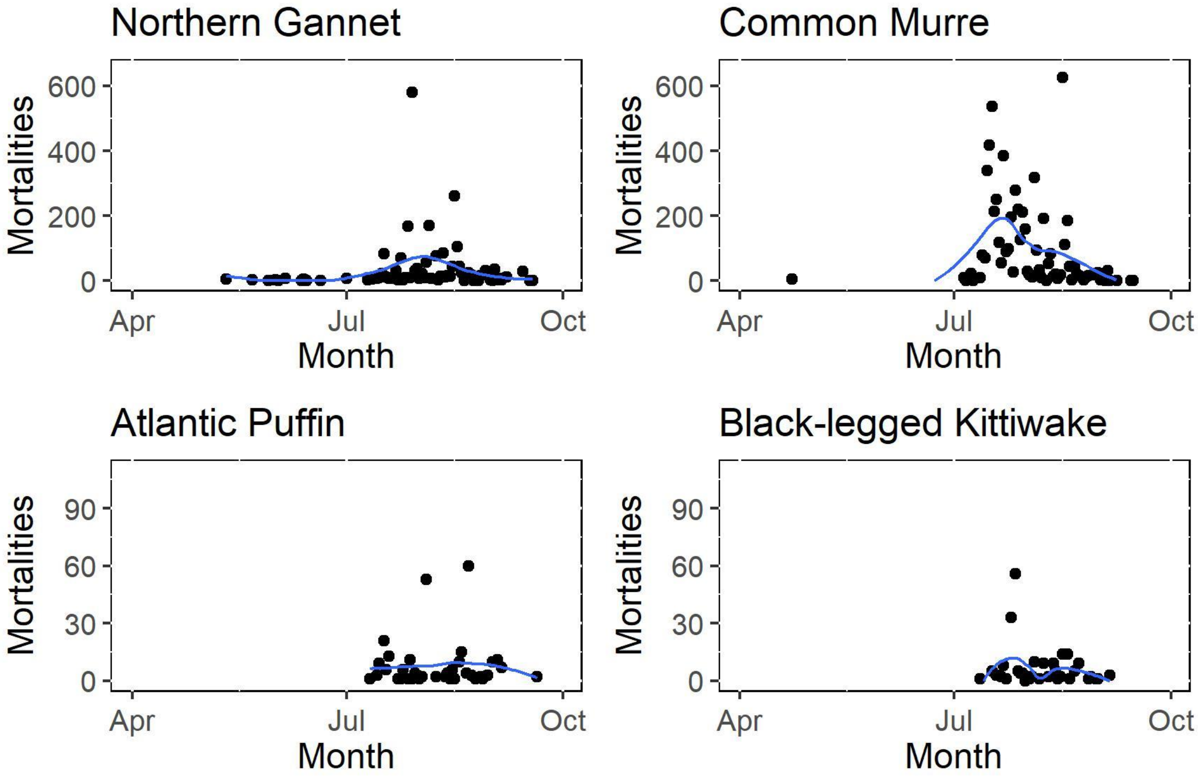


Fig. S3. Frequency of observed mortalities among Northern Gannets, Common Murres, Atlantic Puffins, and Black-legged Kittiwakes on the island of Newfoundland from May to September 2022. Note the different y-axis scale for Atlantic Puffins and Black-legged Kittiwakes.

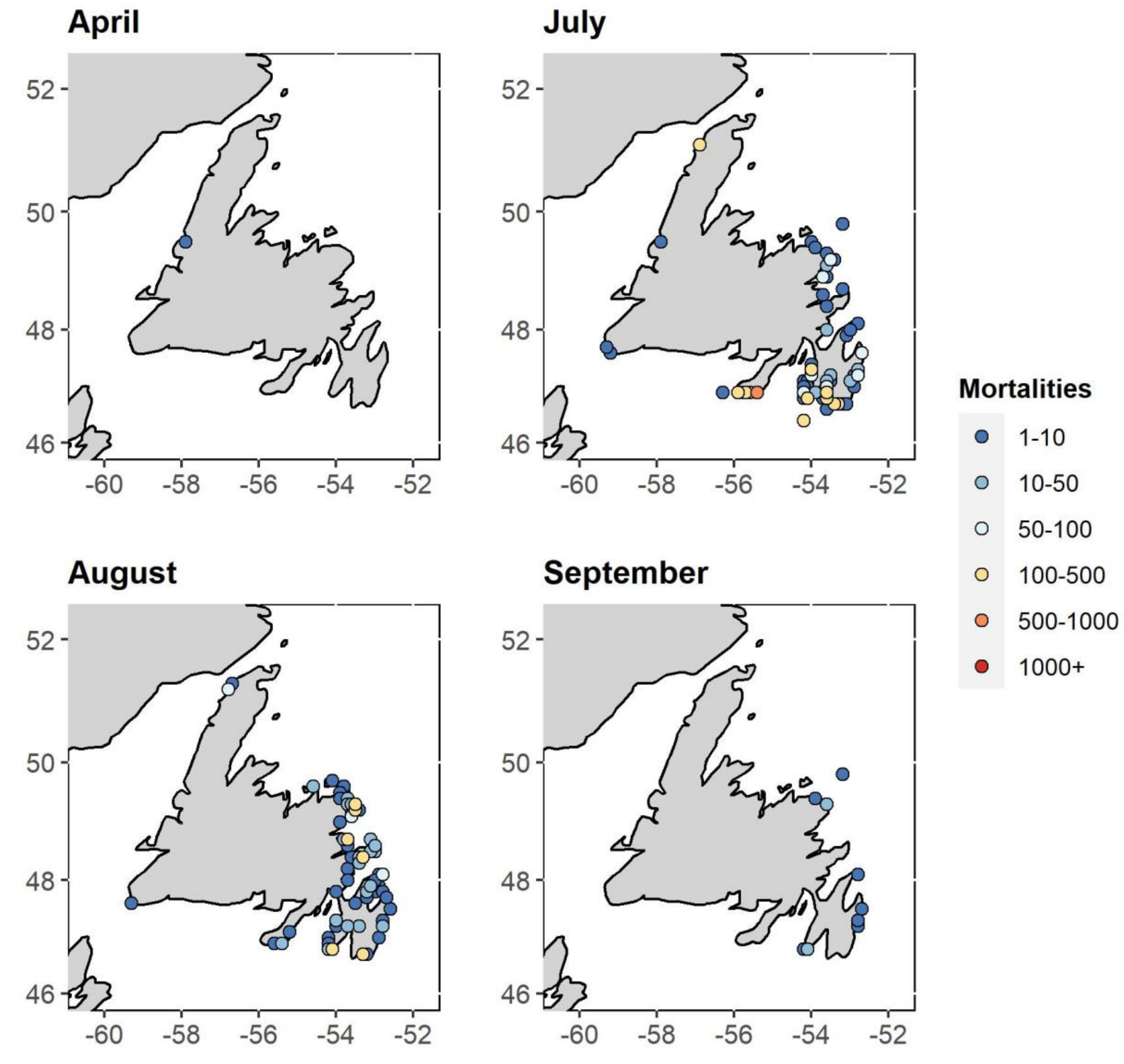


Fig. S4. Common Murre monthly mortality associated with HPAI H5N1 on the island of Newfoundland.

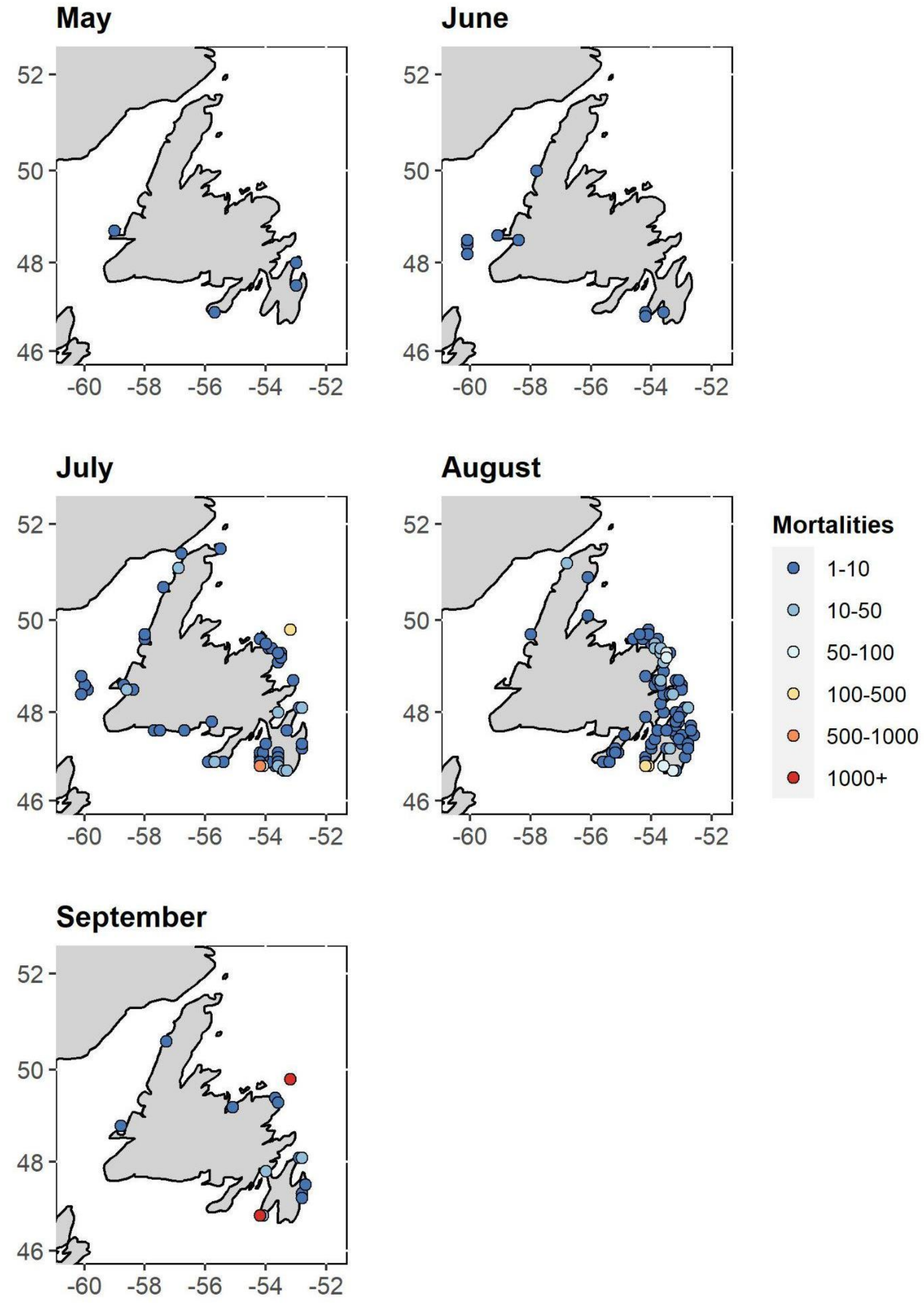


Fig. S5. Northern Gannet monthly mortality associated with HPAI H5N1 on the island of Newfoundland.

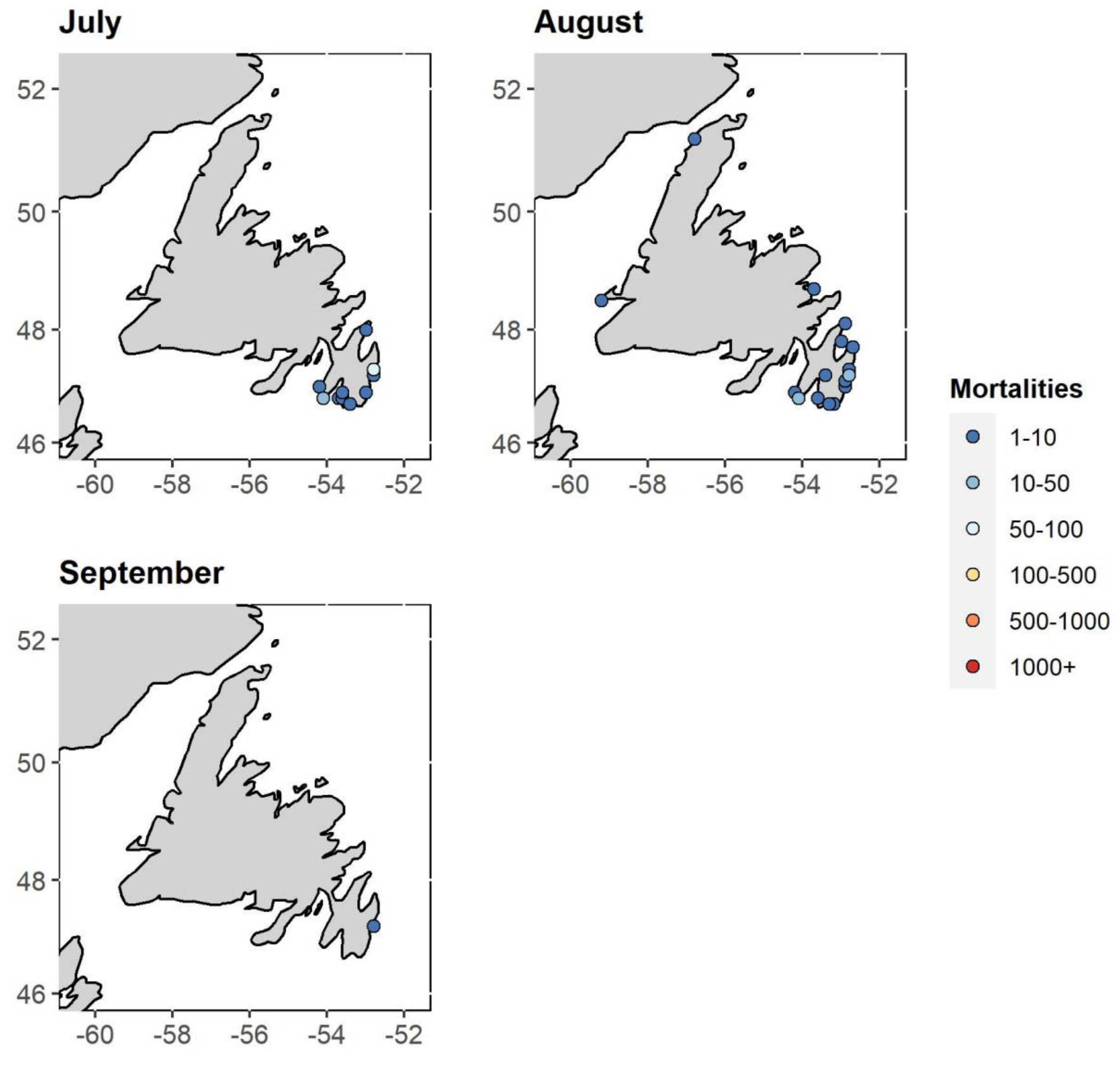


Fig. S6. Black-legged Kittiwake monthly mortality associated with HPAI H5N1 on the island of Newfoundland.

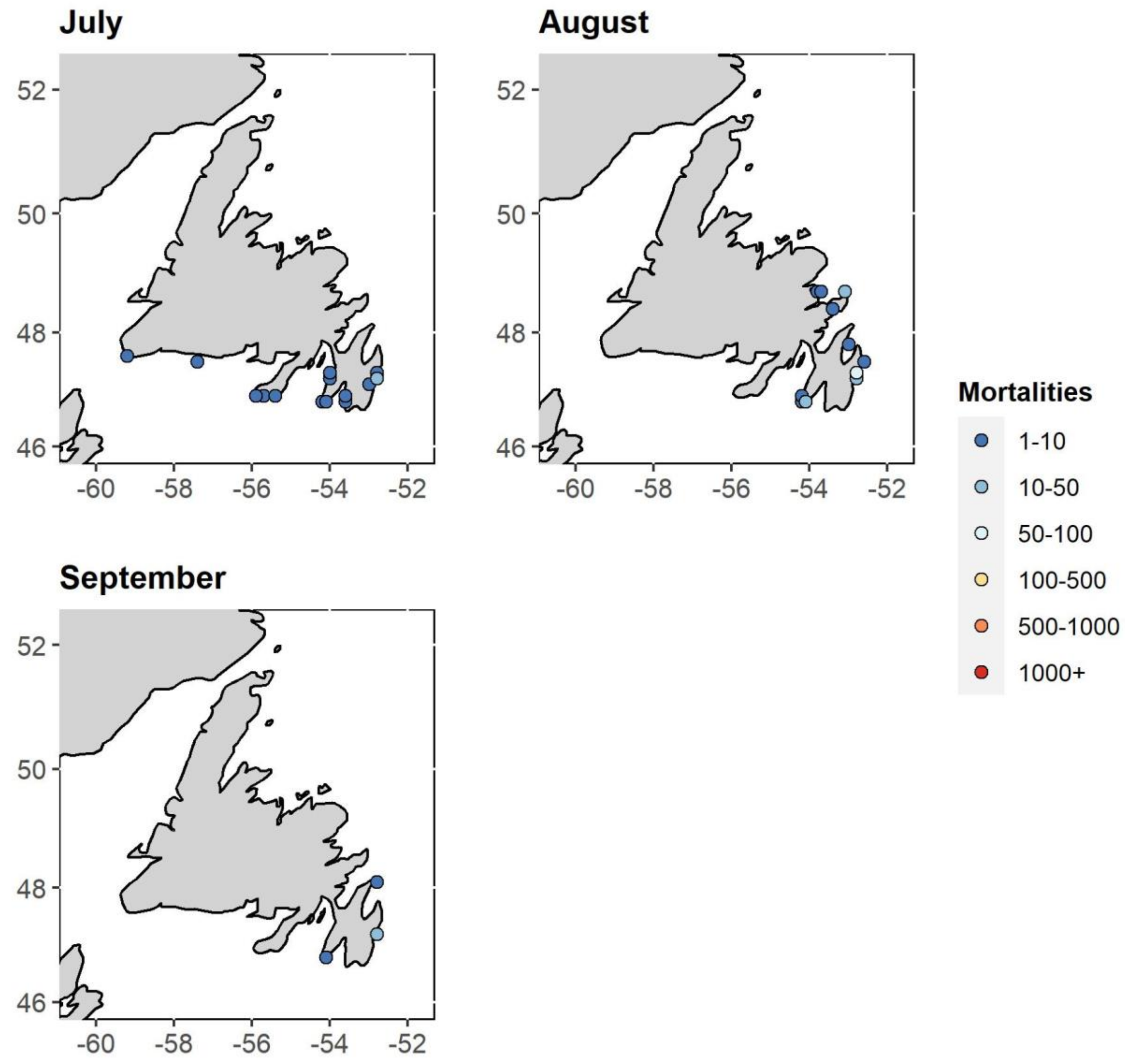


Fig. S7. Atlantic Puffin monthly mortality associated with HPAI H5N1 on the island of Newfoundland.

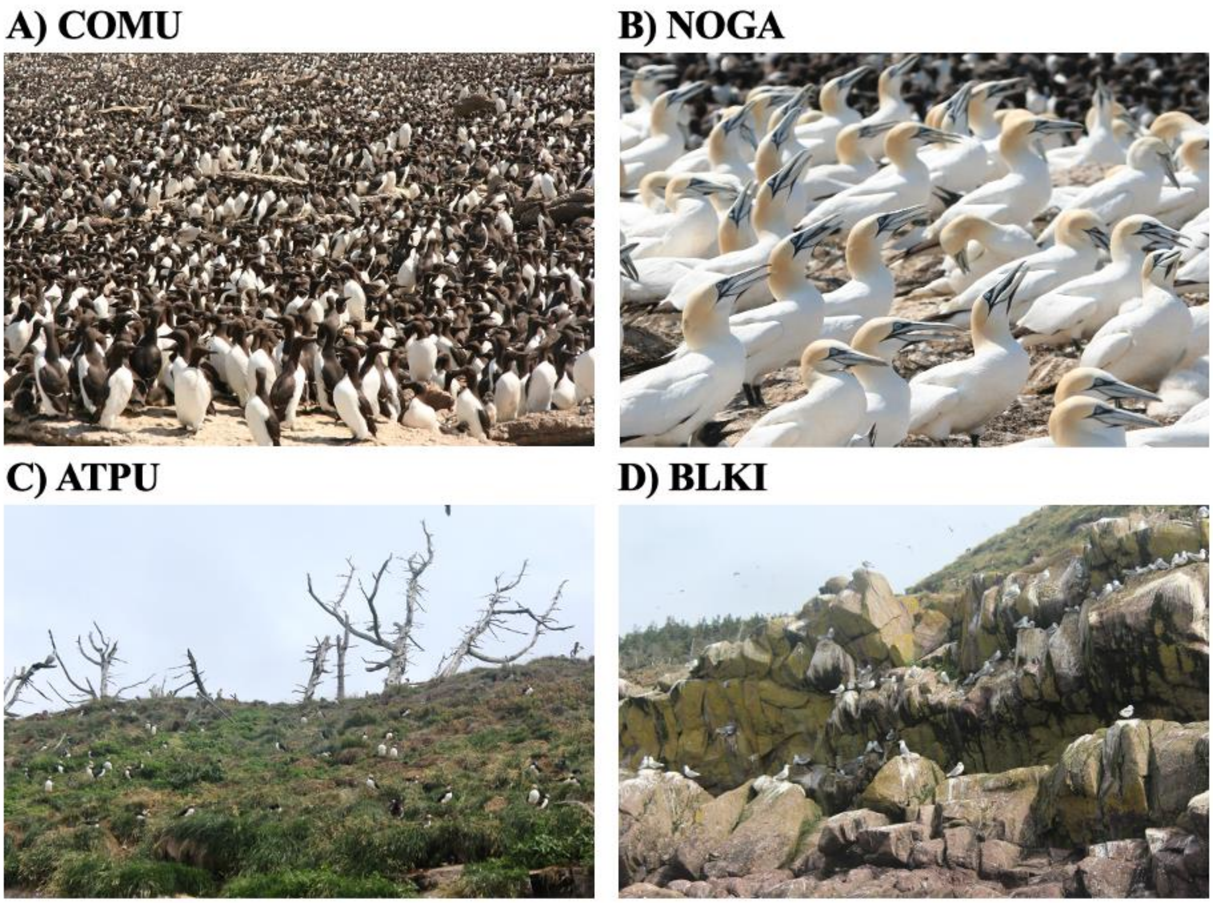


Fig. S8. Breeding aggregation of (A) Common Murres on Funk Island, eastern Newfoundland [Bill Montevecchi]; (B) Northern Gannets on Bonaventure Island [Bill Montevecchi]; (C) Atlantic Puffins on Gull Island, Witless Bay [Gretchen McPhail]; (D) Black-legged Kittiwakes on Gull Island, Witless Bay [Gretchen McPhail].

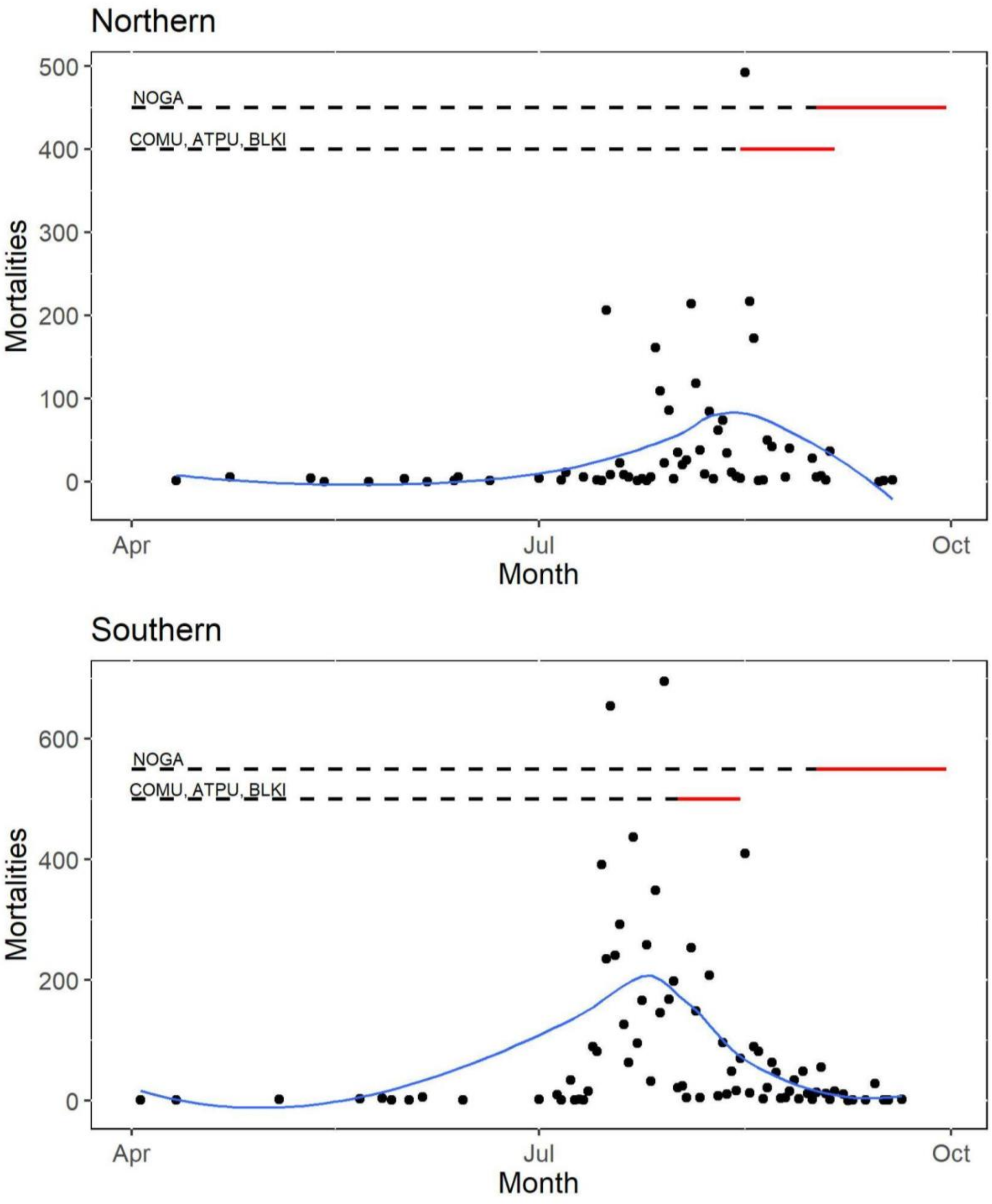


Fig. S9. Breeding season (dashed horizontal line) and colony departure period (red horizontal line) of Northern Gannets (NOGA), Common Murres (COMU), Atlantic Puffins (ATPU), and Black-legged Kittiwakes (BLKI) compared to the daily sum of reported mortalities (black points) for the Northern and Southern region of the island of Newfoundland, Canada. The blue line is the LOESS line of smoothing. Northern is here defined as any location north of 48.1° latitude (the most northern point of the Avalon Peninsula).
