## Supplementary figures and images for "Geographic, ecological, and temporal patterns of seabird mortality during the 2022 HPAI H5N1 outbreak on the island of Newfoundland"

### Fig. S2

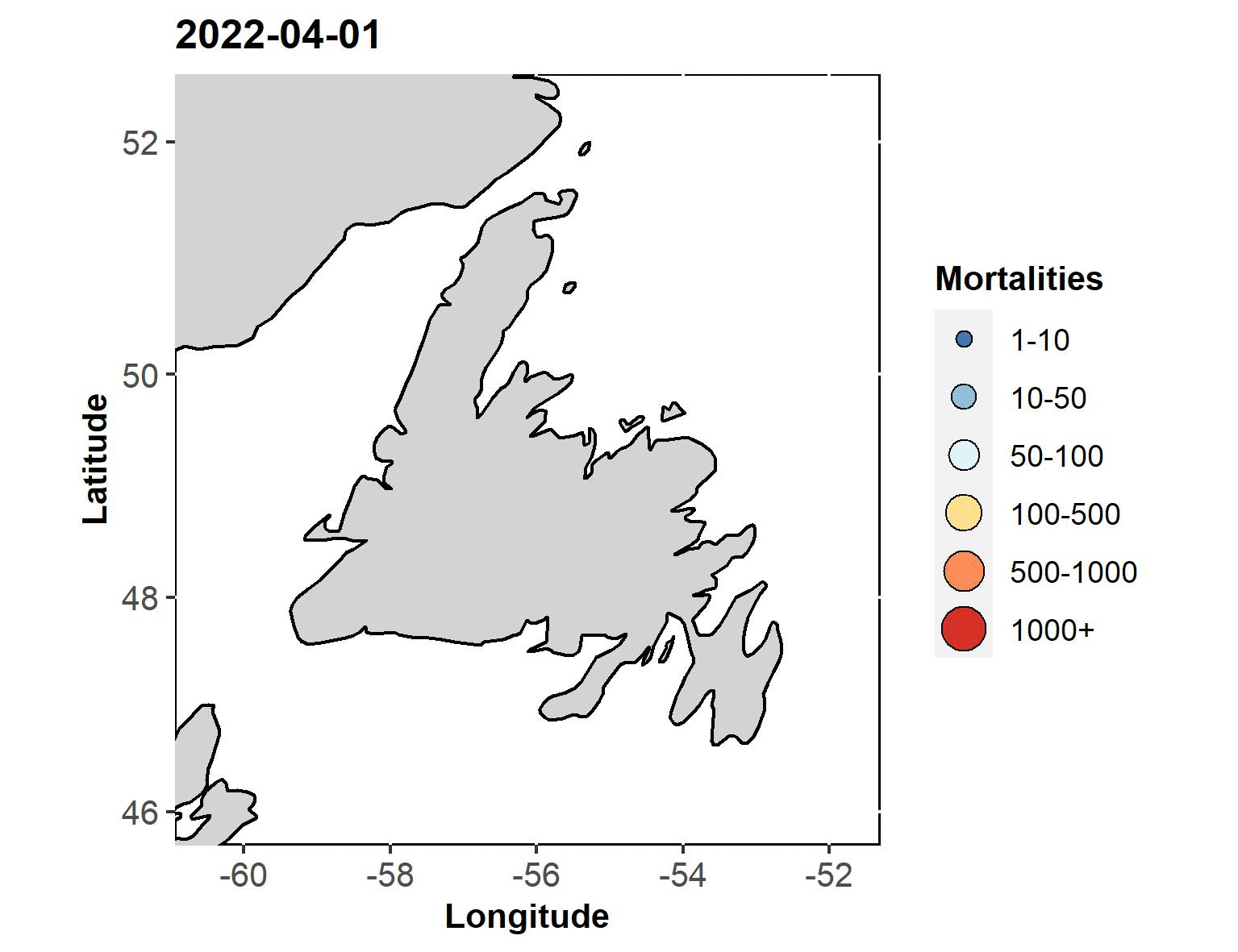
